## Supplementary material for "Data-driven learning how oncogenic gene expression locally alters heterocellular networks": PeerReviewToDate_NatureBiomedicalEngineering

The following Supplement contains the peer-review documents associated with the submission of the pre-print:

To Nature Biomedical Engineering. Items include:

- Initial decision letter.
- Rebuttal letter 1.
- Decision letter and associated peer-review (major revision)
- Rebuttal letter 2.

**Subject:** Decision on Article nBME-20-1233
**Date:** Wednesday, May 20, 2020 at 9:31:35 AM Eastern Daylight Time
**From:** Nature Biomedical Engineering
**To:** David Klinke

Dear Prof Klinke,

Thank you for submitting to *Nature Biomedical Engineering* your Article, "Data-driven learning how oncogenic gene expression locally alters heterocellular networks". I regret that we are unable to consider it for external peer review.

As you may know, we screen manuscripts against our editorial criteria, and decline a substantial proportion of them without input from external referees. In such cases, even if reviewers were to certify the manuscript as technically correct, we feel that it would not be of outstanding interest to merit publication in *Nature Biomedical Engineering*. These editorial judgements are based on considerations of the degree of advance, broad implications, and breadth and depth of the work reported in the manuscript.

The topic of the manuscript is within the remit of the journal. However, in view of your published work on identifying causal networks linking cancer processes and anti-tumor immunity using Bayesian network inference and metagene constructs, in addition to your work covering an unsupervised strategy for identifying epithelial-mesenchymal transition state metrics in breast cancer and melanoma, we feel that your data-driven approach, using digital cytometry and Bayesian Network inference, to link increases in oncogenic gene expression with alterations in stromal and immune cell subsets from bulk transcriptomic data does not reach the high threshold CHOICE_null_in_degree_of_advance_in_breadth_and_depth_in_broad_implications that we look for in comparable manuscripts that we consider for external peer review.

Nevertheless, the work may be appropriate for *Nature Communications*, to which it could be transferred by using the link in the footnote below.

Please be assured that this editorial decision does not represent a criticism of the technical quality of the work. We hope that you will soon be able to see the manuscript published.

Best wishes,

Rosy

__
Dr Rosy Favicchio
Senior Editor, *Nature Biomedical Engineering*

* If you wish to transfer your manuscript to *Nature Communications*, please use our manuscript transfer portal to initiate the transfer to this journal (or to another journal of your choice published by Nature Research). This transfer link remains active until used. If you transfer your manuscript to Nature-branded journals or to the Communications journals, you will not have to re-supply manuscript metadata and files.

All Nature Research journals are editorially independent, and the decision to consider manuscripts are taken by their own editorial staff. For more information, please see our manuscript transfer FAQ page.

This email has been sent through the *Nature Biomedical Engineering* Springer Nature Manuscript Tracking System.

*Confidentiality Statement: This e-mail is confidential and subject to copyright. Any unauthorized use or disclosure of its contents is prohibited. If you have received this email in error please notify our helpdesk.*

Confidentiality and pre-publicity policy | Privacy Policy | Update Profile

**Subject:** Re: Decision on Article nBME-20-1233
**Date:** Wednesday, May 20, 2020 at 2:52:12 PM Eastern Daylight Time
**From:** David Klinke
**To:** Nature Biomedical Engineering
**CC:**

Hi Dr. Favicchio,

Thank you for your response and I'd have to say I'm really surprised (the cynic in me not so much). While I know that it is a judgement call involving:
- **the degree of advance:** we posit that the work could realize a rule-based modeling approach for cell-level networks, which doesn't currently exist. Current methods rely on math models hand curated by experts.
- **broad implications:** we're focusing on immuno-oncology, which in 2019 became the leading category of pharmaceutical agents in RnD and is one of the most competitive. Mechanistic math modeling and simulation is becoming increasingly important in pharma RnD (see NIH white paper https://www.nigms.nih.gov/training/documents/systemspharmawpsorger2011.pdf ). So much so that Certara created a consortium with abbvie, astellas, Bayer, Merck, Shire and Takeda to create – by hand curation - a immuno-oncology simulation platform. https://www.certara.com/pressreleases/certara-launches-industry-first-quantitative-systems-pharmacology-qsp-consortium-on-immuno-oncology-with-leading-pharma-company-members/?ap=Array&UTM_LeadSource= Their initial effort created an ensemble model from existing models https://www.certara.com/wp-content/uploads/2019/09/Chelliah_2019_PAGE_immunotherapies.pdf . One of the take home points from our work is the importance of tumor cell functional plasticity and fibroblasts – neither are represented in this ensemble model. That helps drive home the point related to the problem with hand curated models – they only include what experts think are important instead of letting the data speak for themselves.
**- breadth and depth of the work reported:** This is an integrated computational and experimental study that builds off of prior work. As you know, no paper is written in a vacuum. In the email, two other papers are mentioned, which I have highlighted in yellow and green below. Paper 1 is a computational study with no experimental validation and uses an unrefined approach for defining cellular features (i.e., a hand curated) and for network inference (we used only one algorithm). Paper 2 is also a computational study that lays some groundwork in defining cellular features (in a machine learning sense) for the functional plasticity of malignant cells within an epithelial to mesenchymal state space. Of note, functional plasticity was represented as a lumped feature and was confounded with fibroblasts in Paper 1, which are criticisms that we addressed in Paper 2. I know that the manuscript is a bit long – but we have to cover a lot of ground here as the study combines computational and experimental aims. CCN4 is also a novel mechanism by which tumor cells can promote local immunosuppression, which is the focus of the included supplementary manuscript.

In light of those comments, would you consider this manuscript for external peer review?
Sincerely,
David Klinke

—
David J. Klinke II, Ph.D.
Professor - Department of Chemical and Biomedical Engineering
Adjunct Associate Professor - Department of Microbiology, Immunology & Cell Biology
West Virginia University, Morgantown, WV 26506-6102
(304) 293-9346 | ESB 517 | BMRF 219 | Website: http://klinke.openwetware.org

**From:** Nature Biomedical Engineering <>
**Reply-To:** Nature Biomedical Engineering <>
**Date:** Wednesday, May 20, 2020 at 9:31 AM
**To:** David Klinke <>
**Subject:** Decision on Article nBME-20-1233

Dear Prof Klinke,

Thank you for submitting to *Nature Biomedical Engineering* your Article, "Data-driven learning how oncogenic gene expression locally alters heterocellular networks". I regret that we are unable to consider it for external peer review.

As you may know, we screen manuscripts against our editorial criteria, and decline a substantial proportion of them without input from external referees. In such cases, even if reviewers were to certify the manuscript as technically correct, we feel that it would not be of outstanding interest to merit publication in *Nature Biomedical Engineering*. These editorial judgements are based on considerations of the degree of advance, broad implications, and breadth and depth of the work reported in the manuscript.

The topic of the manuscript is within the remit of the journal. However, in view of your published work on identifying causal networks linking cancer processes and anti-tumor immunity using Bayesian network inference and metagene constructs <Paper 1>, in addition to your work covering an unsupervised strategy for identifying epithelial-mesenchymal transition state metrics in breast cancer and melanoma <Paper 2>, we feel that your data-driven approach, using digital cytometry and Bayesian Network inference, to link increases in oncogenic gene expression with alterations in stromal and immune cell subsets from bulk transcriptomic data does not reach the high threshold CHOICE_null_in_degree_of_advance_in_breadth_and_depth_in_broad_implications that we look for in comparable manuscripts that we consider for external peer review.

Nevertheless, the work may be appropriate for *Nature Communications*, to which it could be transferred by using the link in the footnote below.

Please be assured that this editorial decision does not represent a criticism of the technical quality of the work. We hope that you will soon be able to see the manuscript published.

Best wishes,

Rosy

__
Dr Rosy Favicchio
Senior Editor, *Nature Biomedical Engineering*

* If you wish to transfer your manuscript to *Nature Communications*, please use our manuscript transfer portal to initiate the transfer to this journal (or to another journal of your choice published by Nature Research). This transfer link remains active until used. If you transfer your manuscript to Nature-branded

journals or to the Communications journals, you will not have to re-supply manuscript metadata and files.

All Nature Research journals are editorially independent, and the decision to consider manuscripts are taken by their own editorial staff. For more information, please see our manuscript transfer FAQ page.

This email has been sent through the *Nature Biomedical Engineering* Springer Nature Manuscript Tracking System.

*Confidentiality Statement: This e-mail is confidential and subject to copyright. Any unauthorized use or disclosure of its contents is prohibited. If you have received this email in error please notify our helpdesk.*

Confidentiality and pre-publicity policy | Privacy Policy | Update Profile

**Subject:** Request for the reporting summary and policy checklist for Article NBME-20-1233A-Z
**Date:** Friday, May 22, 2020 at 11:06:22 AM Eastern Daylight Time
**From:**
**To:** David Klinke
**CC:**

Dear Prof Klinke,

Thank you for submitting additional clarifications in regards to the advances made in your Article, "Data-driven learning how oncogenic gene expression locally alters heterocellular networks". Having re-assessed the work, we have decided to send the manuscript out to external review. However, before we contact potential reviewers, I should ask you to please fill in our reporting summary and policy checklist. (Please note that the reporting summary will be available to the reviewers, and these forms are dynamic PDF files that can only be properly visualized and filled in by using Acrobat Reader.)

Both documents are aimed at ensuring good reporting standards as well as compliance with policies on research ethics. Should the manuscript be published, the reporting summary will be attached to the PDF version of the paper and will also be available as supplementary information. More information is available on the editorial policies page.

Also, please update the submitted manuscript files should you need to modify the manuscript to include any missing information on statistics or methods or to update the plots to conform to the recommended data-presentation policies.

When you have completed both the reporting summary and the checklist, please upload both forms as well as any updated manuscript files. (*The link in this paragraph will allow you to securely send the files to us via Dropbox. You do not need to have a Dropbox account. Please make sure that each filename follows the pattern ####CorrespondingAuthorSurname_FileDescription, where #### are the four last digits of the manuscript tracking number, and note that we will get automatically notified when your files have been uploaded.*)

Best wishes,

Rosy

—
Dr Rosy Favicchio
Senior Editor, *Nature Biomedical Engineering*

This email has been sent through the *Nature Biomedical Engineering* Springer Nature Manuscript Tracking System.

*Confidentiality Statement: This e-mail is confidential and subject to copyright. Any unauthorized use or disclosure of its contents is prohibited. If you have received this email in error please notify our helpdesk.*

Confidentiality and pre-publicity policy | Privacy Policy | Update Profile

**Subject:** Decision on Article NBME-20-1233A-Z
**Date:** Thursday, August 6, 2020 at 8:55:56 AM Eastern Daylight Time
**From:** Nature Biomedical Engineering
**To:** David Klinke

Dear Professor Klinke,

Thank you again for submitting to *Nature Biomedical Engineering* your manuscript, "Data-driven learning how oncogenic gene expression locally alters heterocellular networks" and please excuse the unusual delay in reaching a decision. The manuscript has been seen by three experts, whose reports you will find at the end of this message and that, despite our chasing efforts, one reviewer did not provide a report. You will see that although the reviewers have some good words for the work, they articulate concerns about the degree of support for the claims, and in this regard provide useful suggestions for improvement. We hope that with significant further work you can address the criticisms and convince the reviewers of the merits of the study. In particular, we would expect that a revised version of the manuscript provides:

* improvements to the methodological section and a more robust justification for the design of the model as a data-driven analytical tool.

* clarifications as to the extent of prior knowledge and assumptions required for network inference.

When you are ready to resubmit your manuscript, please upload the revised files, a point-by-point rebuttal to the comments from all reviewers, the (revised, if needed) reporting summary, and a cover letter that explains the main improvements included in the revision and responds to any points highlighted in this decision.

Please follow the following recommendations:

* Clearly highlight any amendments to the text and figures to help the reviewers and editors find and understand the changes (yet keep in mind that excessive marking can hinder readability).

* If you and your co-authors disagree with a criticism, provide the arguments to the reviewer (optionally, indicate the relevant points in the cover letter).

* If a criticism or suggestion is not addressed, please indicate so in the rebuttal to the reviewer comments and explain the reason(s).

* Consider including responses to any criticisms raised by more than one reviewer at the beginning of the rebuttal, in a section addressed to all reviewers.

* The rebuttal should include the reviewer comments in point-by-point format (please note that we provide all reviewers will the reports as they appear at the end of this message).

* Provide the rebuttal to the reviewer comments and the cover letter as separate files.

We hope that you will be able to resubmit the manuscript within 25 weeks from the receipt of this message. If this is the case, you will be protected against potential scooping. Otherwise, we will be happy to consider a revised manuscript as long as the significance of the work is not compromised by work published elsewhere or accepted for publication at *Nature Biomedical Engineering*. Because of the COVID-19 pandemic, should you be unable to carry out experimental work in the near future we advise that you reply to this message with a revision plan in the form of a preliminary point-by-point rebuttal to the comments from all reviewers that also includes a response to any points highlighted in this decision. We should then be able to provide you with additional feedback.

We hope that you will find the referee reports helpful when revising the work. Please do not hesitate to contact me should you have any questions.

Best wishes,

Rosy

—
Dr Rosy Favicchio
Senior Editor, *Nature Biomedical Engineering*

---

Reviewer #1 (Report for the authors (Required)):

Klinke and colleagues set out to develop mechanistic models that link different cell types via their expression profiles. From this they propose it is possible to identify causal relationships, including some involved in oncogenesis. There is a lot to like about this research and the work: it combines a range of state-of-the-art theoretical and computational methods, it makes concrete and biologically relevant predictions which are then tested.

I believe that the introduction and concluding section are clear and accessible, the results — which include the computational methodology — are really quite involved and accessibility will almost certainly depend on the readers' backgrounds; clearly the work appears to be of uniformly high standard, but some of the sections that are outside my area were hard-going and required a fair bit: terminology should be defined more consistently and with the intended broad audience in mind.

My detailed comments will therefore focus on those bits of the paper that are closest to my own research. In general the work flow in Fig 1 serves as a nice framework to understand the general direction of the research. I would have liked to see more detail on the mapping of genes-patients and genes-cell-types to cell-types-patients and how this is conducted in practice.

The main issues where I would like to have clarification, beyond this, relate to the reconstruction of the Bayesian network.

The structure learning seems to involve a number of steps that could be rationalised more clearly. First, the consensus seed network reconstruction relies on an ensemble of networks and while the description flows logically, there are a number of questions remaining: is this done in the context of the methods outlined in reference 26? The selection of methods would otherwise need more detail. The value of ensemble estimators has rarely been rigorously tested and one estimator is included, ARACNE, that have come to find extremely unreliable: the false positives generated by this algorithm really cause a problem in all the instances that I am familiar. In the same vein the use of correlations to assign edge directions is based on quite severe assumptions (e.g. there are problems that there is no easy way to distinguish between direct and indirect interactions in several of the methods).

Then there are assumptions in including certain edges (Lines 152-160), which are only partially justified (I trust that the authors can do so easily), but here and at a later stage (Lines 198-204) it would be interesting to see how such choices affect/guide/bias the network construction.

Such limitations of the search space are, of course, very helpful, but it is not clear as to how easily these methods are applied to new data or in new contexts. Is domain expertise a limiting factor? There is no problem if that were the case (anyhow essential for meaningful analysis and interpretations of results), but it should be stated explicitly, somewhere.

On lines 247-248 I am unsure as why are these values chosen? Is there a way to make this more a priori rigorous? For example, looking at network inference as a multiple testing problem could be appropriate?
One question also related to this regards the choice of CCN4 an example: is this based on the authors' previous experience, and expertimental expertise? Or is it a system that is particularly amenable to this analysis (low-hanging fruit)?

Assessing the validation section becomes easier once the questions above have been addressed. As it stands the analysis appears appropriate, but it is unclear as to whether the chosen interventions are maximally informative. The statistical analysis conducted here needs to include multiple testing corrections. I fear that upon introduction of FDR or similar techniques, many of the "significant" results (eg. p=0.047, p=0.022) will vanish. I would like to see seed and Bayesian network construction better explained and I think that following that the assessment of the validation will become easier.

Detailed comments:
Figure 2: this figure is important but not very clear. First, the max count is 9 and this would make the presentation cleaner than going up to 10. Why are some of the universally inferred edges excluded? I would like to see a description of edges 21,22,25,26 etc and why they are not included in the network. Understanding this will be important to assess I feel that getting this right will be important to ensure correctness downstream in the workflow.

L48-52: I think that this represents only a subsection of problems confounding data; references 6 and 7 are not capturing the present state very well.

L 147 "with" or "to" missing

L 182 bootstrapping what?

L247-248: why are these values chosen? Is there a way to make this more a priori rigorous? Links to local FDR?

L317-322 direction of edges and presence of edges?

L 450 p-values are odd

L544 p-value now given as p<0.05, Why?

I wave my right to anonymity,
Michael Stumpf

Reviewer #2 (Report for the authors (Required)):

The authors developed a method to analyze cell regulatory network by combining digital cytometry technology with Bayesian network inference, which allows the analysis of cell-to-cell subpopulation regulatory networks from mixed cell population data. The method has a great advantage for reducing the number of nodes in the network by using cell subpopulations rather than using genes in the cell, which make it possible the analysis of cell-cell regulation at the tissue level. Based on the idea, this method also solves one of the problems with Bayesian networks, since a large number of the node (genes) are considered to be a bottleneck to infer the network accurately. This method can estimate the network in a data driven way (cell population data in this study), as opposed to mechanic mathematical models that require deep biological knowledge and hand curation. Current study predicted how increased CCN4 expression might alter the cell-cell regulatory network in oncogenesis of breast
and melanoma patients and validated the prediction using cell culture and mouse models.

Overall comments
This method is unique in that it can analyze cell-to-cell regulation at the cell population level. It would be useful to directly analyze the efficacy of immune cell therapy against cancer using single-call and bulk sequence data, predict the time trajectory of cell population changes to lead diagnosis at early stage of the treatment. This method has the potential for such medical applications. One of the drawbacks of this current method is that it still requires substantial hand curations before actually inferring the network. For example, CCN4 is specified and some cell-cell interactions are preconfigured. Without this kind of information, it seems difficult to analyze the number of nodes used in the network. We understand that patient numbers and variation noise affect the accuracy of inference results and consistent predictions. However, in a real world, single cell sequencing can find more than just the number of cell populations in the tissue. This problem has to be solved
and it has to be in a more data-driven way if the authors intend to develop an analytical method to utilize digital cytometry for diagnosis or test the efficacy of drug treatment or cell therapies.

Major comments

- Digital cytometry method developed by Newman et al. (Nature Biotech 2019) appeared to be powerful in distinguishing cell subpopulations from a mixed population and the effects of genetic mutations on cell population components and prognosis. To take full advantage of the information available from digital cytometry, current Bayesian method may not be a good way. In addition, current study uses an already existing R package tool. Therefore there is no novelty here in the Bayesian method.

- But the number of nodes may vary between patients based on the cell subpopulations separated by digital cytometry. So how the results would be changed if digital cytometry classifies different cell populations (nodes) from patient to patient? For example, patient A has 7 nodes and patient B has 9 nodes. Even in this case, important core regulatory relationships are either held constant or it predicts totally different regulation.

Minor comments
- Please explain the "blacklist" and "whitelist" more clearly. What are the benefits of using these lists for inference?

- Please describe the biological function of coherent feedforward motifs in networks. How this network is associated with cancer malignancy in terms of cell-cell interactions?

Reviewer #3 (Report for the authors (Required)):

Klinke et al. propose an approach to network inference at the cell-level that aims at reconstructing the causal relationships between genetic drivers of oncogenesis and other phenotypic changes due to cancer progression that might be derived from transcriptomic datasets. The network inference is unsupervised (apart from the incorporation of some prior knowledge), relies on 9 existing algorithms but it is combined to digital cytometry to make the network lower-dimensional. The causal relationships inferred are compared to mouse models for melanoma.

I think that the problem per se of network reconstruction from these datasets at the cell-level is extremely relevant, given the important contribution of the tissue micro-environment to cancer evolution. The degree of advance provided by this work is essentially methodological, however not fully original - the original contribution is the the formulation of the network at the cell-level using ‘features’ from digital cytometry while I understand that ref 26 essentially provides the algorithmic machinery. In addition, I did not get the impression that the steps followed were meant to provide a methodological pipeline that could be more broadly applied. I think that the lack of clarity and completeness about these steps contributed substantially to this impression.
To be more concrete, I have 2 major reservations against the publication of this manuscript (unless it undergoes revisions), which are the following:
1) Many details of the algortihmic procedure are missing or stated without precision, which can give the impression

that the steps followed were not rigorous. Major technical criticisms or questions in this regard:
- Lines 149-165: The description of the construction of the network can be improved. In particular: I would add a list, or at least examples, of the 'features' included: I would explain how 'root' - 'leaf' nodes are defined, based on the connectivity; I would say clearly what are the criteria to create the 'blacklist' (on the latter, the sentence at line 210, mentioning 'prior knowledge' is too vague). In particular, it is not clear how the blacklist is used in the inference, given that bayesan inference has a mathematical way to include prior knowledge by assigning a prior probability to each edge (or generally each parameter to learn).
Furthermore, understanding how 'features' are chosen would also help better understand the sentence at line 488, mentioning the uncertainty about features in myeloid subsets in mouse models.
- Line 170: I would make clearer the connection between the choice of the 9 algorithms and ref 26. If they are not a prescription of the method in ref 26, they all need a separate reference and a short commetn on why they were chosen.
- Line 185: the number 0.5 is hard to understand without a comment on how the algorithm work (so what this number represents), maybe this could be included in the methods.
- Line 187: I would need to know how the correlation coefficient is calculated, maybe this could be included in the methods.
- Lines 215 and 708: It is not clear how the maximum likelihood estimation of parameters from a Gaussian ansatz was formulated.
- Line 695: How this transformation to numerical values was made needs a clarification.

2) I find difficult from the discussion and the figures to understand the evidence upon which the statement about the consistency between inferred network predictions and the experimental validation (line 550) is based. Here I think that making the reader able to extract this information from the figures is essential, which would help also understand what results in particular prove, in a statistically significant way, that the model is predictive.
Major technical criticism or questions in this regard:
- Fig. 4: What is the connection between normalized CCN4 and the the conditional probability model? Does the correlation coefficient refer to the orange or the blue linear fit? What correlation in particular confirms in a clearcut way the inference predictions?
- Fig. 6 and 7: Here it would be helpful if some visual element could be used to connect these results to the topology of the network inferred when the results are used to corroborate the network predictions.

Minor comments:
- Line 21: In the abstract I think it should be mentioned what is the result of the tests made.
- Line 45: I would insert refs. where digital cytometry and bayesan inference are first introduced;
- Line 103: I would stress that the small number of biological replicates not only makes the task difficult computationally, but also implies a greater uncertainity in the inference;
- Line 149: I think one needs to say what the learning algorithms used are, if these are the same 9 metioned below then I would add 'see below';
- Fig 2: How was the edge strength estimated and how is its trend on the figure explained?

This email has been sent through the *Nature Biomedical Engineering* Springer Nature Manuscript Tracking System.

*Confidentiality Statement: This e-mail is confidential and subject to copyright. Any unauthorized use or disclosure of its contents is prohibited. If you have received this email in error please notify our helpdesk.*

Confidentiality and pre-publicity policy | Privacy Policy | Update Profile

**Response to Reviewer's Comments**

**Data-driven learning how oncogenic gene expression locally alters heterocellular networks.**

David J. Klinke II, Audry Fernandez, Wentao Deng, Habibolla Latifizade, and Anika C. Pirkey

Submitted for review in Nature Biomedical Engineering (Manuscript ID: NBME-20-1233A-Z)

Overall, we thank the reviewers for their service and their constructive criticisms on the manuscript. We feel that this revised paper presents, in a more comprehensive way, our findings related to inferring how genetic alterations associated with oncogenesis alters heterocellular networks within tissues. In particular, the critiques identified:

- a need for more robust justification for the design of the model as a data-driven analytical tool

In response to this critique, key revisions include a more detailed explanation of the "community network" approach using a blacklist that incorporates known biological constraints as to how information can flow within the heterocellular network and incorporation of a Bayesian Information Criterion (BIC) applied to the individual nodes to justify including arcs in a whitelist. As a more formal approach, the BIC helps infer a network structure that is parsimonious for each node, that is it includes enough arcs to capture the dependence among nodes but not too many that limits interpretability.

- clarification as to the extent of prior knowledge and assumptions required for network inference

In response to this critique, key revisions include a more detailed explanation of the rationale for including arcs in a blacklist and results that illustrate the impact of including particular arcs in the blacklist. As discussed in more detail below, we believe that these changes enhance the data-driven nature of the approach.

Our responses to the specific comments are listed below in blue font. A red font is used to indicate the text that has been changed in this revised manuscript.

Peer Reviewer #1 Evaluations:

Klinke and colleagues set out to develop mechanistic models that link different cell types via their expression profiles. From this they propose it is possible to identify causal relationships, including some involved in oncogenesis. There is a lot to like about this research and the work: it combines a range of state-of-the-art theoretical and computational methods, it makes concrete and biologically relevant predictions which are then tested.

I believe that the introduction and concluding section are clear and accessible, the results — which include the computational methodology — are really quite involved and accessibility will almost certainly depend on the readers' backgrounds; clearly the work appears to be of uniformly high standard, but some of the sections that are outside my area were hard-going and required a fair bit: terminology should be defined more consistently and with the intended broad audience in mind.

My detailed comments will therefore focus on those bits of the paper that are closest to my own research. In general the work flow in Fig 1 serves as a nice framework to understand the general direction of the research. I would have liked to see more detail on the mapping of genes-patients and genes-cell-types to cell-types-patients and how this is conducted in practice.

The main issues where I would like to have clarification, beyond this, relate to the reconstruction of the Bayesian network.

1. The structure learning seems to involve a number of steps that could be rationalised more clearly. First, the consensus seed network reconstruction relies on an ensemble of networks and while the description flows logically, there are a number of questions remaining: is this done in the context of the methods outlined in reference 26? The selection of methods would otherwise need more detail. The value of ensemble estimators has rarely been rigorously tested and one estimator is included, ARACNE, that have come to find extremely unreliable: the false positives generated by this algorithm really cause a problem in all the instances that I am familiar. In the same vein the use of correlations to assign edge directions is based on quite severe assumptions (e.g. there are problems that there is no easy way to distinguish between direct and indirect interactions in several of the methods).
   Authors Response (AR): Reference 26 provides a convenient software interface to generate comparable results while swapping out network inference algorithms. The ensemble approach is not described in Ref 26 and follows from the "community network" approach similarly described by Marbach and coworkers based on experiences with DREAM challenges (Marbach et al Nature Methods (2012)9:796, Marbach Prill et al. PNAS 2010) A "community prediction" combines network predictions obtained by multiple inference methods and performs better than any individual algorithm. Marbach et al. also mention that community performance is robust to including an algorithm that performs poorly - as suggested for ARACNE. We note though that we have removed ARACNE in the revised version. Yes, two of the methods included in the revised version (MMPC and HITON) do not provide directed edges. We included these algorithms in the Community prediction stage to include additional evidence supporting the existence of an edge between two nodes.

2. Then there are assumptions in including certain edges (Lines 152-160), which are only partially justified (I trust that the authors can do so easily), but here and at a later stage (Lines 198-204) it would be interesting to see how such choices affect/guide/bias the network construction.
   AR: We denoted the "Cancer" node as a root node as genetic alterations within a particular cell type, such as a melanocyte in the case of melanoma, can transform a normal tissue into a malignant tissue. CCN4 was also denoted as a child of "Cancer" but as a potential root node for all other nodes based on evidence that CCN4 is produced by

malignant cells (Deng JBC 2019, Deng CMB 2020), at a higher level in malignancy (Deng JBC 2019, Klinke PLoSCompBio 2014), and by an intracellular regulatory network that appears to be dysregulated in malignant cells (Deng JBC 2019, Klinke MBoC 2015). CD8 T cells were denoted as a leaf node as they are a key cell type that can elicit targeted cytotoxicity to control tumor growth. We have included additional text to clarify these choices. We have also included results where we progressively impose these assumptions in the blacklist (new Fig. S1).

3. Such limitations of the search space are, of course, very helpful, but it is not clear as to how easily these methods are applied to new data or in new contexts. Is domain expertise a limiting factor? There is no problem if that were the case (anyhow essential for meaningful analysis and interpretations of results), but it should be stated explicitly, somewhere.
   AR: Limiting the network search space is done using a blacklist in generating a community edge list and both a blacklist and whitelist in inferring the network structure used in estimating the model parameters. In formulating a blacklist, specifying cancer as a root node and CD8 T cells as a leaf should be generalizable in the context of most tumors. Due to immune privilege, glioblastoma may be an exception for CD8 T cells (Kaiser et al. Biotech Prog 2016). Specifying the oncogene as a child of cancer and root node to everything else should also be generalizable. Single cell RNAseq data can help support this assumption that malignant cells express the oncogene. Specifying nodes that have a lot of zero values as leaves follows from network inference arguments as trends in the non-zero values, which ideally are related to other nodes, may be swamped by differences between the zero and non-zero values, which may give rise to spurious arcs. We have now included text to state this explicitly.

4. On lines 247-248 I am unsure as why are these values chosen? Is there a way to make this more a priori rigorous? For example, looking at network inference as a multiple testing problem could be appropriate?
   AR: Lines 247-248 relate to queries of the joint conditional probability distribution and implications of representing a discrete variable as a continuous variable. Specifically, the "Cancer" node only has values of 0 (a normal sample) or 1 (a sample taken from tumor tissue). However, the joint conditional probability distribution is represented by a linear Gaussian model where the Cancer node is assumed to be a continuous variable. The conditional probability distribution is queried by generating random samples and retaining samples consistent with the query. The probabilities are defined for a range of values, i.e., "Cancer" < 0.15 as the probability of sampling a discrete value, i.e., Cancer = 0, would be zero. Tightening the range, say "Cancer" < 0.05, doesn't appreciably change the figures but increases the number of samples required to generate the figure as more are rejected. In this revision, we have redone the analysis with "Cancer" < 0.05 and "Cancer" > 0.95. In terms of structural learning, we use a resampling approach to generate 10,000 network structures for each algorithm and then use the average network structure. This resampling approach improves the rigor of the structural learning step.

5. One question also related to this regards the choice of CCN4 an example: is this based on the authors' previous experience, and experimental expertise? Or is it a system that is particularly amenable to this analysis (low-hanging fruit)?
   AR: The focus on CCN4 (aka WISP1) is based on the authors' previous experience. We identified it using a proteomics screen for tumor-secreted factors that suppress immune response to Interleukin-12 in 2012 (DOI: 10.1039/c2ib20053h ). CCN4 is not particularly well studied (only 469 papers are listed in PubMed searching for CCN4/WISP1/ELM1). In DREAM network inference challenges, investigators are typically blinded to the ground truth. Outside of a DREAM challenge, blinding is more difficult when collective knowledge about a network is high. Here, collective knowledge about CCN4 is relatively sparse and translation of knowledge acquired in the mouse to human systems is an important concern. We have acquired experimental data to validate aspects of the predicted network. The network inference results also motivate additional studies to clarify aspects of the network, like the interaction between CCN4 and cancer associated fibroblasts. We obtained some interesting preliminary results but the COVID-19 slowdown limits our ability to pursue those experiments. Overall, we felt that this data-driven approach for inferring heterocellular networks could be applied to other contexts.

6. Assessing the validation section becomes easier once the questions above have been addressed. As it stands the analysis appears appropriate, but it is unclear as to whether the chosen interventions are maximally informative. The statistical analysis conducted here needs to include multiple testing corrections. I fear that upon introduction of FDR or similar techniques, many of the "significant" results (eg. p=0.047, p=0.022) will vanish. I would like to see seed and Bayesian network construction better explained and I think that following that the assessment of the validation will become easier.
   AR: Now, we have revised our discussion of how we seed and construct the Bayesian networks and how the results are validated. In particular, we have focused on how the digital cytometry results are validated and how the Bayesian Networks are validated. Of note, using a FDR approach is not used conventionally for analyzing flow cytometry data as the gating strategy makes each measurement not purely independent, which is why we have included supplemental figures illustrating the gating. We would also say that showing p-values is intended to quantify the strength of evidence rather than for explicit hypothesis testing.

   Detailed comments:

7. Figure 2: this figure is important but not very clear. First, the max count is 9 and this would make the presentation cleaner than going up to 10. Why are some of the universally inferred edges excluded?
   AR: We have updated this figure, as mentioned above, to reflect now 10 structural learning algorithms, the impact of the whitelist threshold and node-based BIC that are used to generate a consensus seed list. As mentioned in the revised version to motivate the BIC, including arcs in a network is a trade-off between capturing the joint probability

distribution and interpretability of the network. Some of the universally inferred edges are excluded as including them may lead to overfitting (i.e., they are not at the minimum BIC) or that they may create cycles.

8. I would like to see a description of edges 21,22,25,26 etc and why they are not included in the network. Understanding this will be important to assess I feel that getting this right will be important to ensure correctness downstream in the workflow.
   AR: We have also modified the labeling scheme so that the edge numbers are conserved across all three data sets. In the revised manuscript, the equivalent edges in the BRCA analysis are 18, 24, 25, 28, 29, and 30. Arc 18 is Epithelial -> Neutrophils, which is excluded as it has a high strength (6E-4) and the minimum BIC value for Neutrophils is with no parental nodes. Arc 24 is pM2 -> Neutrophils, which has a lower strength than Arc 18 (7E-9) but the minimum BIC value for Neutrophils is with no parental nodes. Arc 25 is Macrophages -> Neutrophils, which is also excluded because of the BIC value. Arc 29 is active NK cells -> CD4 T cells, which has a high strength (1E-3) and the minimum BIC occurs at a Whitelist threshold of 1E-5. The same criteria applies to Arc 30 : pM1 -> CD4 T cells, which has a strength of 1E-2.

9. L48-52: I think that this represents only a subsection of problems confounding data; references 6 and 7 are not capturing the present state very well.
   AR: In the interest of brevity, we unfortunately can't review the relevant literature. We hope that the citations provide the reader with some leads to that body of work. If the reviewer feels that one or two other references better capture the present state, we would be happy to change the citations.

10. L 147 "with" or "to" missing
    AR: Thanks for the keen eye, have now included "with".

11. L 182 bootstrapping what?
    AR: We have now included a brief explanation of what was bootstrap resampled.

12. L247-248: why are these values chosen? Is there a way to make this more a priori rigorous? Links to local FDR?
    AR: We discuss in point 4 above. In brief, we have redone the analysis with "Cancer" < 0.05 and "Cancer" > 0.95. A linear Gaussian model is used to generate samples. These values are just used to filter samples consistent with the discrete "Cancer" variable (i.e., Cancer = 0 corresponds to data from a normal tissue sample while Cancer = 1 corresponds to data obtained from a cancer tissue sample).

13. L317-322 direction of edges and presence of edges?
    AR: We have revised the text for clarity - edges do not have direction while arcs refer to a direction of influence.

14. L 450 p-values are odd

AR: We have redone panel 7D as some of the data points that were used in the statistical calculation were missing from the figure.

15. L544 p-value now given as $p<0.05$, Why?
    AR: This has been revised to reflect the actual p-values.

I wave my right to anonymity,

Michael Stumpf

Reviewer #2 (Remarks to the Author):

The authors developed a method to analyze cell regulatory network by combining digital cytometry technology with Bayesian network inference, which allows the analysis of cell-to-cell subpopulation regulatory networks from mixed cell population data. The method has a great advantage for reducing the number of nodes in the network by using cell subpopulations rather than using genes in the cell, which make it possible the analysis of cell-cell regulation at the tissue level. Based on the idea, this method also solves one of the problems with Bayesian networks, since a large number of the node (genes) are considered to be a bottleneck to infer the network accurately. This method can estimate the network in a data driven way (cell population data in this study), as opposed to mechanic mathematical models that require deep biological knowledge and hand curation. Current study predicted how increased CCN4 expression might alter the cell-cell regulatory network in oncogenesis of breast and melanoma patients and validated the prediction using cell culture and mouse models.

Overall comments
This method is unique in that it can analyze cell-to-cell regulation at the cell population level. It would be useful to directly analyze the efficacy of immune cell therapy against cancer using single-call and bulk sequence data, predict the time trajectory of cell population changes to lead diagnosis at early stage of the treatment. This method has the potential for such medical applications. One of the drawbacks of this current method is that it still requires substantial hand curations before actually inferring the network. For example, CCN4 is specified and some cell-cell interactions are preconfigured. Without this kind of information, it seems difficult to analyze the number of nodes used in the network. We understand that patient numbers and variation noise affect the accuracy of inference results and consistent predictions. However, in a real world, single cell sequencing can find more than just the number of cell populations in the tissue. This problem has to be solved and it has to be in a more data-driven way if the authors intend to develop an analytical method to utilize digital cytometry for diagnosis or test the efficacy of drug treatment or cell therapies.
AR: In this revised manuscript, we believe that the approach is more data-driven than the previous iteration. While scRNAseq is a powerful approach to identify different cell types

present within a sample, batch effects are a persistent problem for increasing the power of typical scRNAseq datasets to aid in understanding how oncogenesis alters heterocellular networks. In addition, the number of features tends to be much greater than the number of biological samples. While the problem with batch effects may be solved in the near future, acquiring enough samples to power a study based solely on scRNAseq data will take years if not decades as patient recruitment and standardization among participating centers are key limitations. The scRNAseq issue with statistical power and batch effects is noted in a recent review (Fan, J., Slowikowski, K. & Zhang, F. Single-cell transcriptomics in cancer: computational challenges and opportunities. Exp Mol Med (2020). https://doi.org/10.1038/s12276-020-0422-0 ) Of note, the TCGA study was started around 2005, where the data available today reflects over 15 years of work to acquire and process patient samples in a uniform way. While we recognize that there is room for improvement, the work described in the manuscript is an approach to solve this problem by leveraging the data that we currently have.

Major comments

1. Digital cytometry method developed by Newman et al. (Nature Biotech 2019) appeared to be powerful in distinguishing cell subpopulations from a mixed population and the effects of genetic mutations on cell population components and prognosis. To take full advantage of the information available from digital cytometry, current Bayesian method may not be a good way. In addition, current study uses an already existing R package tool. Therefore there is no novelty here in the Bayesian method.
AR: We agree that digital cytometry is a helpful tool to infer cell subsets from a mixed signature. As mentioned in the text, digital cytometry has been primarily used in correlative studies. We feel that the novelty here is in two aspects. First, we feel that integrating digital cytometry and sample features with Bayesian network inference algorithms is novel. A second novel element is the community prediction approach using a blacklist that incorporates known biological constraints as to how information can flow within the heterocellular network and a Bayesian Information Criterion applied to the individual nodes to generate a whitelist. Yes, the ``bnlearn" package is a convenient interface to a variety of structural learning and utilities but using that package alone is not what we are claiming as a novelty. Collectively, we believe the work describes an analogous rule-based modeling approach for cell-level networks as has been developed for molecular-level networks. Given the recent interest in immuno-oncology, this is an especially important area of unmet need.

2. But the number of nodes may vary between patients based on the cell subpopulations separated by digital cytometry. So how the results would be changed if digital cytometry classifies different cell populations (nodes) from patient to patient? For example, patient A has 7 nodes and patient B has 9 nodes. Even in this case, important core regulatory relationships are either held constant or it predicts totally different regulation.
AR: To clarify, digital cytometry estimates changes in abundance of different cell types within a tissue - that is are there more or less CD8+ T cells present within a patient's

tumor tissue. The number of potential nodes of the network is equal to the number of features associated with the dataset. In the absence of time-course data obtained from the same system, we are using the collective dataset obtained for a particular cancer that occurs at the same anatomical location to infer the structure of the network and then the parameters describing the joint probability distribution for feature prevalence is conditioned on the structure of the network. As far as inferring different regulation (i.e., network structure) based on one patient sample, that doesn't really make sense as one can't draw network inferences from n = 1.

Minor comments

3. Please explain the "blacklist" and "whitelist" more clearly. What are the benefits of using these lists for inference?
   AR: In this revised manuscript, we have rewritten our description of how we selected the "blacklist" and "whitelist" to enhance clarity. In short, the benefit of specifying arcs as part of the "blacklist" is that non-physiologic arcs (e.g., an increase in CD8 T cells cause cancer) are not considered for inclusion into the network. Arcs included in a "whitelist" promote the inference of a network structure that is parsimonious for each node, that is it includes enough arcs to capture the dependence among nodes but not too many that limits interpretability.

4. Please describe the biological function of coherent feedforward motifs in networks. How this network is associated with cancer malignancy in terms of cell-cell interactions?
   AR: Feed-forward loops are highly prevalent and well understood as intracellular control mechanisms and have been extensively studied in gene systems within E.coli and yeast. While there are 8 different feed-forward loop motifs, the one described in the current manuscript is a coherent type 1 feed-forward loop (C1-FFL).

   In a C1-FFL, Promoter X directly promotes both Y and Z while Y goes on to further promote Z. This motif is beneficial to the cell because it allows for controlled regulation of the activation and deactivation of Z through a sign-sensitive delay that will have either an AND or an OR configuration. If both X and Y are required for activation of Z, the C1-FFL will lead to a delayed stimulation as Y builds up to the required level, but will result in no delay when stimulation stops (AND motif). If only one of the regulators is required to stimulate Z, the opposite control mechanism will occur, with an immediate response taking place after stimulation followed by a delayed response of Z after stimulation stops (OR motif).

   The OR motif of the C1-FFL has been studied in the context of flagella motor biogenesis genes in E.coli. Upon activation of FlhDC, genes required to build the flagella motor are immediately activated, while the introduction of FliA (which is activated by FlhDC) has an additive effect of the motor gene activation. Upon deactivation of FlhDC, the motor genes remain active for approximately one hour, which is approximately the time needed for biogenesis of a flagella. Mutations to FliA lead to an immediate deactivation of the flagella motor genes after FlhDC is deactivated, implying an OR motif of a C1-FFL.

The presence of this control loop protects the flagella biogenesis process from transient deactivation of FlhDC, allowing the E.coli to escape nonideal survival conditions even in the event that FlhDC signaling is temporarily lost (Alon, U. Network motifs: Theory and experimental approaches. Nature Reviews Genetics (2007). DOI: 10.1038/nrg2102). This protection from transient deactivation is important in many biological processes. In the context of oncogenesis, CCN4, and the mesenchymal cell state in the melanoma data set, it is unclear whether the C1-FFL motif is an AND or OR design and further laboratory work would be needed in the future to fully understand the complexities of this signalling loop.

The motif controlling the mesenchymal cell state inferred by the DAG in the breast cancer data set is a type 3 incoherent feed-forward loop (I3-FFL), which would be an intriguing discovery if proven correct because of the rarity of this loop design in biological processes. However, while the structure of a feed-forward loop is present, the question remains as to whether or not it is truly an I3-FFL motif or whether it may be another motif. These questions are primarily based on the wide range of influence values within the loop. The extent of influence of the negative causal relationship between “cancer” and “mesenchymal” is rather weak (absolute value of 0.17) in comparison to the positive causal relationship between “CCN4” and “mesenchymal”, which is the highest value found in the DAG (absolute value of 1.06). Additional data could ultimately change the type of feed-forward loop that is uncovered and warrants further investigation. Specifically, the GEO data set is a more balanced distribution of benign skin nevi and primary melanoma samples while the BRCA data set contains a lower percentage of normal mammary samples and the tumor samples are all classified as invasive breast cancer, suggesting that an early stage of oncogenesis (i.e., malignant but non-invasive) may be missing.

Feed-forward loops in cell-cell signalling and within cancer have not been well investigated, and laboratory tools to investigate such signalling motifs are more challenging to employ relative to those tools used to study intracellular signalling. Nevertheless, discovery and understanding of these control motifs is an important gap to fill in our understanding as the pharmaceutical industry moves away from gene targeted therapy and towards therapies that modify the cellular interactions within the tumor microenvironment such as tumor vaccines and other immunotherapies.

Reviewer #3 (Remarks to the Author):

Klinke et al. propose an approach to network inference at the cell-level that aims at reconstructing the causal relationships between genetic drivers of oncogenesis and other phenotypic changes due to cancer progression that might be derived from transcriptomic datasets. The network inference is unsupervised (apart from the incorporation of some prior knowledge), relies on 9 existing algorithms but it is combined to digital cytometry to make the network lower-dimensional. The causal relationships inferred are compared to mouse models for melanoma.

I think that the problem per se of network reconstruction from these datasets at the cell-level is extremely relevant, given the important contribution of the tissue micro-environment to cancer evolution. The degree of advance provided by this work is essentially methodological, however not fully original - the original contribution is the formulation of the network at the cell-level using 'features' from digital cytometry while I understand that ref 26 essentially provides the algorithmic machinery. In addition, I did not get the impression that the steps followed were meant to provide a methodological pipeline that could be more broadly applied. I think that the lack of clarity and completeness about these steps contributed substantially to this impression. To be more concrete, I have 2 major reservations against the publication of this manuscript (unless it undergoes revisions), which are the following:

Many details of the algortihmic procedure are missing or stated without precision, which can give the impression that the steps followed were not rigorous. Major technical criticisms or questions in this regard:

1. Lines 149-165: The description of the construction of the network can be improved. In particular: I would add a list, or at least examples, of the 'features' included: I would explain how 'root' - 'leaf' nodes are defined, based on the connectivity; I would say clearly what are the criteria to create the 'blacklist' (on the latter, the sentence at line 210, mentioning 'prior knowledge' is too vague).

AR: We have now revised our description of how we constructed the network in response to this and critiques by the other reviewers.

2. In particular, it is not clear how the blacklist is used in the inference, given that bayesan inference has a mathematical way to include prior knowledge by assigning a prior probability to each edge (or generally each parameter to learn).

AR: In a Bayesian context, a blacklist in structural inference is used to assign the probability to zero of including a specific directed arc. We have added a sentence to clarify this point.

3. Furthermore, understanding how 'features' are chosen would also help better understand the sentence at line 488, mentioning the uncertainty about features in myeloid subsets in mouse models.

AR: In short, each expression signature for a given cell type used in Digital Cytometry corresponds to a feature. These expression signatures can be obtained using different methods - like based on transcriptomics data obtained from single cell RNAseq or bulk RNAseq of purified cell populations. We used the expression signatures - which include specific subsets of genes and their relative expression for each signature - defined in the CIBERSORT (these are based on bulk RNAseq data) and CIBERSORTx (these are based on scRNAseq data) papers. In defining features associated with myeloid cells, we used signatures derived from both single cell RNAseq (the macrophage feature) and bulk RNAseq of purified cell populations (M0, M1, and M2 macrophages features). Of note, M1 and M2 macrophages can be thought of as mature differentiated macrophages while M0 macrophages are committed to macrophage differentiation but not driven to the extreme of M1 and M2 macrophages. Given the overlap in these features, we combined the M0, M1, and M2 features to estimate the functional orientation of the macrophages present. One of the characteristics of the tumor microenvironment is the presence of myeloid-derived suppressor cells (MDSC) (e.g., Marvel and Gabrilovich J. Clin Invest (2015)125(9):3356-64.). MDSCs are morphologically, phenotypically, and functionally heterogenous and result from impaired differentiation of various myeloid precursors into mature myeloid cells, like macrophages, dendritic cells, and neutrophils. Uncertainty in mapping how the gene expression signature in an immature myeloid cell subset overlaps with a differentiated myeloid cell creates uncertainty in predicting cell abundance by digital cytometry. We have added a couple of additional sentences to help clarify this point.

4. Line 170: I would make clearer the connection between the choice of the 9 algorithms and ref 26. If they are not a prescription of the method in ref 26, they all need a separate reference and a short comment on why they were chosen.

AR: The “bnlearn” package is a convenient wrapper to these different network inference algorithms, we have now included references to all 10 algorithms and a short comment as to why they were chosen.

5. Line 185: the number 0.5 is hard to understand without a comment on how the algorithm work (so what this number represents), maybe this could be included in the methods.

AR: This value follows from the bootstrap resampling to infer the network structure, where learning the structure of a network from the data might be different if a slightly different collection of samples were used. In bootstrap resampling, a bootstrap replicate is generated by sampling with replacement from the original data set to create a synthetic dataset that has the same number of samples as the original dataset. A network structure is then learned from this synthetic dataset and stored. This process was repeated 10,000 times to generate a collection of 10,000 DAGs. Next, we calculate the frequency by which each arc appears in this collection. Using a cut-off for the frequency, an ``average” DAG is generated by retaining only those arcs that have a frequency above a threshold, where the value of 0.5 corresponds to an arc being included in at least 50% of the 10,000 DAGs. Instead of specifying an ad-hoc threshold value, we used a statistically based approach described by Scutari and Nagarajan 2013 (PMID: 23395009). We have now included a citation to this paper.

6. Line 187: I would need to know how the correlation coefficient is calculated, maybe this could be included in the methods.

AR: We have changed this sentence to “the sign of the linear coefficient was also used to annotate whether a particular arc promotes or inhibits the target node.”

7. Lines 215 and 708: It is not clear how the maximum likelihood estimation of parameters from a Gaussian ansatz was formulated.

AR: We represent the Bayesian Network (BN) by (G, θ), where G is the DAG over the set of random variables, i.e., features in the observed data. The nodes in G represent random variables, and the edges in G represent the direct dependencies between them. In the BN, each variable is independent of its nondescendents in the graph given the state of its parents, which is consistent with a Markovian property, where the Conditional probability distributions at each node depends only on its parents. The second component $\theta$ denotes the set of parameters, which are regression coefficients for each variable against its parents, in case of Gaussian Bayesian network (A Gaussian Bayesian network is a network in which the distribution of each variable is Gaussian with fixed variance and a mean that is a linear combination of its parents).

The maximum likelihood estimation is aimed to make inferences about the joint probability distribution of the random variables (i.e.; features in the dataset) that are most likely to have generated the data sample. Evaluating the joint density at the observed data sample $D = \{X_1, \cdots, X_N\}$ gives a real-valued function,

$$L(\theta) = p(D|\theta) = \prod_{i=1}^{N} \quad p(X_i|\theta) \qquad (1)$$

which is called the likelihood function and $\theta = \{\theta_1, \cdots, \theta_N\}$ parameters of a model that index the probability distribution of each random variables. Then the likelihood of data (D) being generated from a model with parameters $\theta$ is the product of the probabilities of each example, given the model parameters $\theta$.

For each data example $X_i = \{F_1, \cdots, F_n\}$ where $F_j{}'s$ are the variables (nodes in the BN), relationships between variables that have been captured in the BN structure defined by the DAG enables the Joint probability distributions to be expressed in terms of a product of Conditional probability distributions $p(F_j|U_j, \theta_j)$, where $\theta_j$ is the set of parameters describing the distribution for $F_j$ against its parents $U_j$. Thus,

$$p(X_i|\theta) = \prod_{j=1}^{n} \quad p(F_j|U_j, \theta_j)$$

Then for maximum likelihood estimates of the parameters of the model ($\theta$) that results in giving maximum value of $p(D|\theta)$ we can maximize the $log(p(D|\theta))$ because the natural logarithm $log()$ function is monotonically increasing, therefore;

$$\hat{\theta} = max\ L(\theta) = max\ p(D|\theta) = \ max\ log\ (p(D|\theta))$$

Since in reality our dataset $D$ is a set of data $D = \{X_1, \cdots, X_N\}$, thus by equation (1) we have

$$\hat{\theta} = \ max\ \ log\ log\ \left(p(D|\theta)\right)\ = \sum_{i=1}^{N} \quad log\ (p(X_i|\theta)\ )$$

Taking into account that, when all the variables in a network have linear Gaussian conditional densities, the joint density over $X$ is a multivariate Gaussian [1, 2]. So, by Theorems 7.3 and 7.4 in reference [1], consider the case where all CPDs of variables are linear-Gaussian and for simplicity let's do the process for maximum likelihood estimates of distributions of $F_k$ given its parents $U_k = \{F_1, \cdots, F_{k-1}\}$, therefore

$$p(F_1 = a_1, \cdots, F_{k-1} = a_{k-1}, \theta_k) = \ N(u_k;\ \mu, \sigma^2) \qquad (2)$$

Such that mean of distribution of the node $F_k$ is a linear combination of states of its parent nodes, i.e., $\mu = \beta_0 + \beta_1 a_1 + \cdots + \beta_{k-1} a_{k-1}$, where $\beta_j{}'s$ are the coefficients of a linear regression model of $F_k$ .
To find maximum likelihood estimates, the task is to learn $(k+1)$ parameters in $\theta_k = (\beta_0, \cdots, \beta_{k-1}, \sigma^2)$.
Let's define log- likelihood from Eq. (2) as;

$$log\ log\ (L(\theta_k)) = \sum_{i=1}^{n} -\frac{1}{2} log\ log\ (2\pi\sigma^2) - \frac{1}{2\sigma^2}(\beta_0 + \beta_1 a_1[i] + \cdots + \beta_{k-1} a_{k-1}[i] - a_k[i])^2$$

Take derivative wrt $\beta_0,\ \beta_1, \cdots and\ \beta_{k-1}$ set them equal to zero and solve for the parameters. From the data, substitute values $a_k[i]$, so we will get system of $k$ linear equations and solve them simultaneously. Finally, the same process can be done for finding $\sigma$.

For instance, in the SKCM model, estimating the parameters of the local distributions of the node "**Macrophages (MACs)"**, $\theta_{MACs}$, from Conditional density of (**MACs | CAFs, NK active cells,** $\theta_{MACs}$ ) as $\theta_{MACs} = (\beta_0, \beta_1, \beta_2, \sigma^2) = (-0.017, 0.57, 0.28, 0.042)$ that has been annotated in the DAG of SKCM for **MACs,** which is coefficients for regression model of **MACs** given its parents; **NK active cells** and **CAFs,** such that the mean of **MACs** is as follows;

$$\begin{aligned} \mu_{MACs} &= \beta_0 + \beta_1 * (CAFs) + \beta_2 * (NK\ active\ cells) \\ &= -0.017 + (0.57) * (CAFs) + (0.28) * (\ NK\ active\ cells) \end{aligned}$$

[1] D. Koller, N. Friedman, Probabilistic graphical models. Principles and techniques, MIT Press, 2009.

[2] G. Elidan, I. Nachman, N. Friedman, "Ideal Parent" Structure Learning for Continuous Variable Bayesian Networks, Journal of Machine Learning Research, 8(63):1799–1833, 2007.

8. Line 695: How this transformation to numerical values was made needs a clarification.
   AR: We have added text to clarify this. Specifically, we state that, if the sample was obtained from normal tissue, the value for "Cancer" was set equal to 0; if the sample was obtained from cancer tissue, the value for "Cancer" was set equal to 1.

   I find difficult from the discussion and the figures to understand the evidence upon which the statement about the consistency between inferred network predictions and the experimental validation (line 550) is based. Here I think that making the reader able to extract this information from the figures is essential, which would help also understand what results in particular prove, in a statistically significant way, that the model is predictive.
   AR: To clarify this point, we have added text stating that we focused on two strategies to validate these predictions: using flow cytometric analysis of tumor

infiltrating lymphocytes in syngeneic mouse tumor models to validate the digital cytometry predictions and targeted in vitro experiments to validate the presence or absence of arcs predicted by the Bayesian network inference.

Major technical criticism or questions in this regard:

9. Fig. 4: What is the connection between normalized CCN4 and the conditional probability model?

AR: In short, the point of Figure 4 is to compare the conditional probability model to the underlying digital cytometry data. As Cancer and CCN4 are the key root nodes, the plots show how the abundance of these six cell types varies with CCN4 and Cancer (nominally 0 versus 1). Predictions from the conditional probability model are generated by randomly sampling the Gaussian distributions associated with the relevant nodes and propagating these values using the structure of the network and influence parameters annotated on each arc. As the Cancer node is modeled as a continuous variable with a range from 0 to 1, predictions are filtered such that predictions corresponding to normal tissue are obtained from "cancer" node values between 0 and 0.05 and to cancer tissue are obtained from "cancer" node values between 0.95 and 1. As mentioned in the methods, all of the feature values were log-transformed and then normalized to a dynamic range of 0 - 1.

10. Does the correlation coefficient refer to the orange or the blue linear fit?

AR: We have revised these figures to show the slope for both normal (blue) and cancer tissue (red).

11. What correlation in particular confirms in a clearcut way the inference predictions?

AR: The correlations help validate the digital cytometry predictions.

12. Fig. 6 and 7: Here it would be helpful if some visual element could be used to connect these results to the topology of the network inferred when the results are used to corroborate the network predictions.

AR: We hope that our revisions helped clarify how these experimental results corroborate the network predictions.

Minor comments:

13. Line 21: In the abstract I think it should be mentioned what is the result of the tests made.

AR: In the interest of brevity, we have included a very brief mention of the experimental results in the abstract.

14. Line 45: I would insert refs. where digital cytometry and bayesan inference are first introduced;

AR: We have added a phrase at the end of this sentence to tell the reader that these concepts will be discussed in the following paragraphs.

15. Line 103: I would stress that the small number of biological replicates not only makes the task difficult computationally, but also implies a greater uncertainity in the inference;
    AR: We have included this caveat.

16. Line 149: I think one needs to say what the learning algorithms used are, if these are the same 9 mentioned below then I would add ‘see below’;
    AR: We have added text to state what particular learning algorithms were used and have added text to help guide the reader through the approach.

17. Fig 2: How was the edge strength estimated and how is its trend on the figure explained?
    AR: We have added text to the caption explaining that edge strength corresponds to the probability of a partial correlation between the two nodes of an arc being explained by random chance, given the rest of the network. Specifically, it is the p-value calculated using the exact t test for a Pearson’s correlation coefficient.
